# scPortrait integrates single-cell images into multimodal modeling

**DOI:** 10.1101/2025.09.22.677590

**Authors:** Sophia C. Mädler, Niklas A. Schmacke, Alessandro Palma, Altana Namsaraeva, Ali Oğuz Can, Varvara Varlamova, Lukas Heumos, Mahima Arunkumar, Georg Wallmann, Veit Hornung, Fabian J. Theis, Matthias Mann

## Abstract

Machine learning increasingly uncovers rules of biology directly from data, enabled by large, standardized datasets. Microscopy images provide rich information on cellular architecture and are accessible at scale across biological systems, making them an ideal foundation for modeling cell behavior. However, a standardized image format does not exist at the single-cell level. Here we present scPortrait, an scverse software package for generation, storage, and application of single-cell image datasets. scPortrait reads, stitches and segments raw fields of view with out-of-core computation scaling to larger-than-memory datasets. Parallelization enables rapid extraction of individual cells into a standardized single-cell image format with fast access to accelerate machine learning. scPortrait enables analysis across modalities including images, proteomics and transcriptomics, identifying cancer-associated macrophage subpopulations by morphology and embedding single-cell images into transcriptome atlases. scPortrait turns microscopy images into a reusable resource for integrative cell modeling, establishing single-cell images as a core modality in systems biology.

## Introduction

The advent of machine learning has transformed multiple research fields including natural language processing^1^, computer vision^2–4^ and climate science^5^. Computational models can now detect patterns in complex datasets without external guidance. Given that biological datasets often contain entangled information on multiple underlying processes, and are therefore not straightforward to interpret, applying machine learning to biology research promises to uncover new mechanisms and generate new hypotheses^6,7^.

A limitation of current machine learning approaches is their requirement for large datasets during training^2^. This has hampered their adoption in biology, where data acquisition is often costly and siloed. Fortunately, one of the most readily acquired modalities is imaging, a domain in which machine learning has recently shown tremendous success. Images can be acquired comparatively easily across biological scales, from whole ecosystems to subcellular structures, and they capture information on the spatial and temporal arrangement of a system’s components in exquisite detail^8^. In cell biology, comprehensive image collections now describe cellular architecture^9^, tissue structure^10,11^ and perturbation responses^12^. In addition to increases in scale, recent advances have enabled the joint acquisition of images and other modalities such as genetic information^13,14,^ protein abundances and transcriptomics^15^. Techniques like deep visual proteomics^16^ and spatial transcriptomics^17^ even enable paired collection of images and other modalities directly from tissue samples. The combination of spatially resolved imaging with the complementary and orthogonal molecular information from other modalities makes the resulting datasets a rich substrate for machine learning models^18^.

Recent approaches to building comprehensive models of cellular activity, also called foundation models or virtual cells, critically depend on learning from multiple modalities^6,18,19^. Realizing the potential of the spatial resolution provided by images requires machine learning-compatible data structures that can support integration and large-scale modeling^20^. Storing data at the level of individual cells, the smallest functional units of life, is the de-facto standard in other modalities such as transcriptomics. However, existing image analysis software such as SPACEc^21^, MCMICRO^22^, spatiomic^23^, QuPath^24^, and CellProfiler^25,26^ does not generate single-cell image datasets but instead runs analyses to generate and save collections of features. OME-NGFF, a recently proposed storage format for biological images, currently does not provide a specification for saving single-cell images^27^. In addition, it saves images as collections of individual files in the zarr format, slowing down random access required when training machine learning models.

To address these limitations we present scPortrait, a software package and file format to process and store single-cell images (https://github.com/MannLabs/scPortrait). Through efficient parallelization and out-of-core computation, scPortrait accelerates the generation of standardized image datasets for machine learning from raw microscopy images on high-performance compute clusters. The resulting .h5sc files enable fast random access, reproducibility and integration with the scverse ecosystem, establishing images as a first-class modality for machine learning. We demonstrate the power of scPortrait for cross-modality modeling by annotating fluorescently imaged tonsillitis samples with gene expression data generated *in silico* using flow matching^28–30^ and by embedding single cells into transcriptome atlases based on their images. We also use morphological information to identify a tumor-associated macrophage subset. We expect scPortrait to become an integral part of future efforts to model cell behavior across modalities.

## Results

### scPortrait generates single-cell image datasets at scale

scPortrait (https://github.com/MannLabs/scPortrait) is a software package and file format that transforms raw microscopy into standardized, analysis-ready single-cell images (Fig. 1). It ingests data from common sources, such as TIFF and zarr files, stitches individual fields-of-view^31^, applies segmentation with built-in or external algorithms^32^ and extracts individual cells into single-cell image datasets (Methods, Fig. S1a). These functions can be run in batches or included in workflow managing systems such as snakemake^33^ or nextflow^34^, using out-of-core computation to deal with larger-than-memory input images. Its open design allows individual processing steps in scPortrait to be run with integrated algorithms or to be offloaded to external pipelines. All intermediate results are saved as SpatialData^35^ objects that can be inspected interactively^36^. CPU- and GPU parallelization speeds up processing. Using 64 threads, scPortrait extracts more than 700 cells per second (Fig. S1b, c). Extracted images are saved in our newly defined .h5sc file. The .h5sc format stores single-cell images based on the HDF5 containers of AnnData^37^ and the scverse ecosystem^38^, ensuring compatibility with existing and future tools^35,39–45^. By enabling fast reading and writing, the .h5sc format accelerates modern machine learning applications (Fig. S1d).

**Figure 1.**
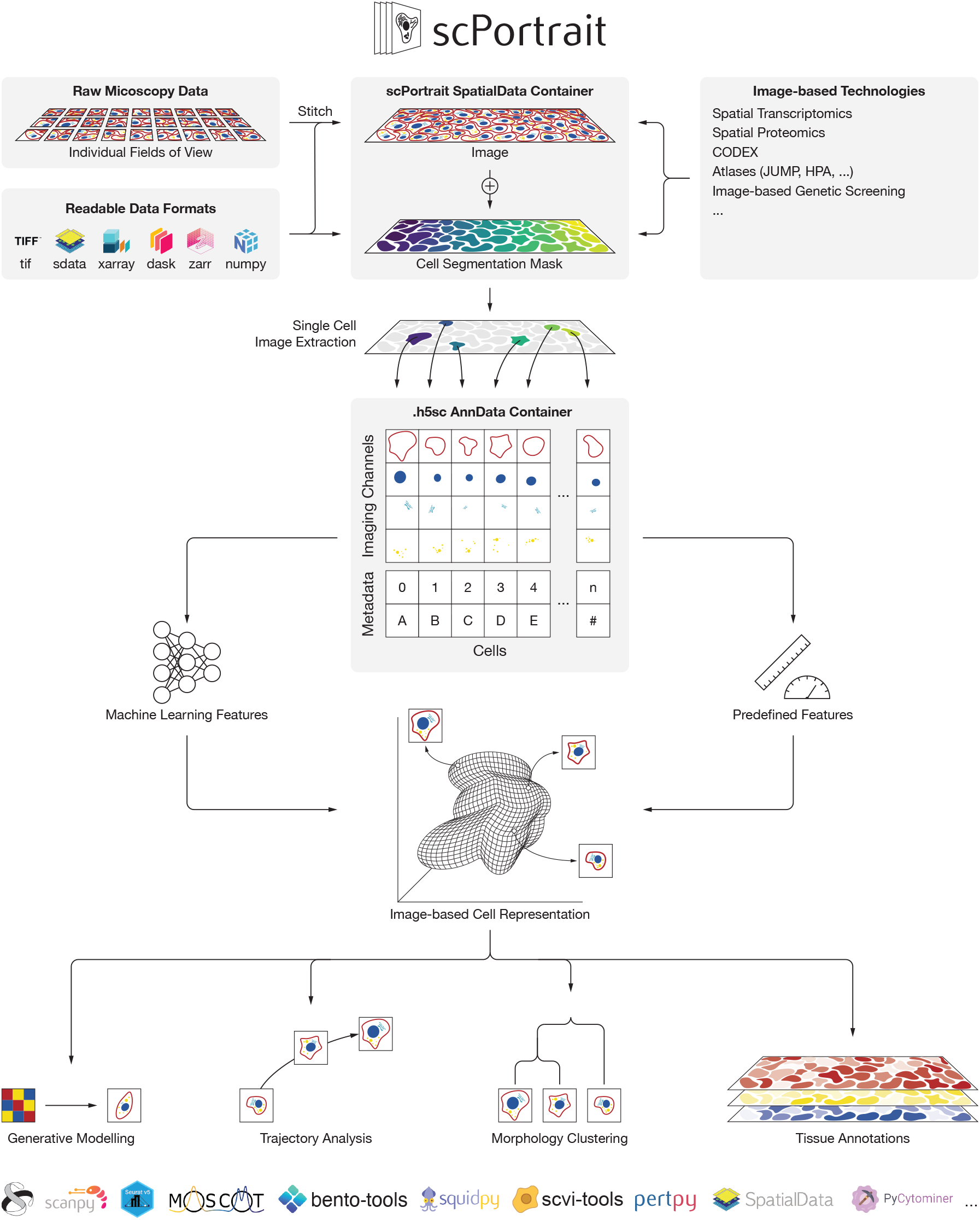
The scPortrait software package and file format for single-cell image dataset generation. scPortrait generates single-cell image datasets from raw imaging inputs. It reads common image formats and stitches raw fields-of-view with high precision. Segmentation masks can then be created with built-in algorithms, external tools, or loaded directly. All intermediates are saved as SpatialData objects for compatibility with external annotations and third-party software. scPortrait then produces standardized single-cell datasets by applying segmentation masks for extraction of single-cell images. Out-of-core computation handles larger-than-memory datasets and all steps are parallelized to enable rapid processing. The resulting single-cell image datasets are stored in an HDF5-based .h5sc file format built on AnnData. This format is directly compatible with downstream analysis steps such as representation learning and integrates with existing software tools for single-cell analysis including scanpy and the scverse ecosystem.

As we demonstrate below, scPortrait powers analyses built on generative modeling and morphology assessment and maps inferred cellular attributes back into the spatial domain, for example into tissue contexts. By making single-cell image storage findable, accessible, inter-operable, and reusable (FAIR)^46^, scPortrait provides the foundation for cross-dataset and cross-modality cellular representation learning, powering analyses from generative modeling to morphology-based tissue mapping^47^.

To demonstrate that scPortrait supports diverse single-cell image embedding strategies, we generated a Golgi morphology benchmark using reporter HeLa cells expressing a fluorescent trans-Golgi marker. We then stimulated these Golgi reporter cells with compounds known to affect Golgi organization and imaged their Golgi morphology (Fig. S2a). Following processing with scPortrait, we featurized the resulting single-cell images in three ways: with ConvNeXt^48^, a convolutional neural network (CNN) model pre-trained on natural images, with SubCell^49^, a transformer-based model pre-trained on cellular images of the Human Protein Atlas^9^ and with CellProfiler^25,26^, a software that extracts pre-defined cellular features (Fig. S2b, c). Despite their different architectures, all embedding strategies clustered cells according to treatment, while separating them from untreated controls (Fig. S2c). In line with previous efforts to extract phenotypic information from image embedders^50^, this shows that all perturbations induce reproducible and separable Golgi morphologies (Fig. S2a, c). We release this Golgi morphology dataset with scPortrait as a ready-to-use benchmark for comparing image-embedding methods.

### Tissue modeling across modalities with scPortrait

scPortrait has already proven invaluable in the analysis of 120 million single-cell images from 40 million cells in a genome-scale image-based genetic screen for autophagosome formation^14^ and in classifying the distinct intracellular distributions of the protein α1-antitrypsin in patient samples of the liver disease α1-antitrypsin deficiency (AATD)^51^. To highlight how scPortrait enables inference across modalities in a disease context, we analyzed a 59-plex CODEX imaging dataset of human tonsils affected by tonsillitis^21^, which contained 1.1 million images of almost 20,000 cells after segmentation (Fig. S3a, b, c). To explore how individual cells contribute to tissue architecture beyond what imaging alone can provide, we sought to map publicly available dissociated CITE-seq data of human tonsils^52^ onto this CODEX dataset. Since these datasets originate from different samples, there is no mapping between individual cells across modalities and neither dataset can be expected to contain all cell types or -states. This makes cross-modality alignment a non-trivial challenge, ideally suited to scPortrait’s integrative framework.

To address this challenge, we turned to optimal transport, a mathematical framework that determines the most efficient way to map one distribution to another. In biology, optimal transport has recently been used to construct developmental trajectories over time and in space^42,53^. Because CODEX, via antibody staining and imaging, and CITE-seq, via sequencing of antibody-conjugated nucleotides, both measure protein abundances, we reasoned that these data should be well suited to generating a probability matrix linking cells across modalities under optimal transport constraints. Using this mapping, we then predicted gene expression based on CODEX features. However, this mapping is discrete, potentially suffering from sampling bias, and it does not necessarily contain all cell states of interest. To generalize beyond observed cell pairs and continuously infer gene expression based on CODEX features, we turned to flow matching^29,30^. This framework constructs probability paths that transform noise into data based on optimal transport maps, making it an ideal fit for our image-to-gene expression modeling problem. Flow matching has previously been used to predict cell behavior in response to diverse stimuli and perturbations^54–56^. To generate gene expression features conditioned on CODEX features, we trained our flow matching model by sampling pairs of source (CODEX) and target (gene expression) cells according to the probabilities in our optimal transport map (Fig. S3d).

To evaluate whether the gene expression profiles inferred by flow matching retained biological information, we tested for recovery of canonical cell type markers. We assigned cell types by k-nearest neighbors prediction to generated gene expression profiles in CITE-seq space. Indeed, we found enrichment of classic markers such as LYZ in monocytes and dendritic cells, CD3 in T cells and NKG7 in NK cells (Fig. S3e). As an aggregate measure of flow matching accuracy, UMAP representations of the measured and flow matching-inferred gene expression spaces overlap substantially, despite expected differences in cell type proportions between CODEX and CITE-seq data (Fig. S3f). These data show that flow matching predicts plausible gene expression features conditioned on CODEX data.

We then used our flow matching model to infer the expression of *TCL1A*, a marker gene of germinal center B cells (GCBC) that was not measured in the CODEX dataset. The inferred profiles revealed clusters of *TCL1A*-expressing cells precisely localized to germinal centers (Fig. S3g, h). Similarly, when inferring the expression of T cell marker *CD2*, we find strong colocalization with CD3 in the tonsil tissue (Fig. S3i, j). Beyond single marker genes, optimal transport enables transfer of higher-level annotations such as cell types from the CITE-seq reference onto the tissue, revealing structures including germinal centers (Fig. S3k). These data demonstrate that image-based cross-modality modeling can recover missing molecular features and reconstruct tissue organization, even when samples across modalities are not matched.

### Modeling morphology of cells in tissues with scPortrait

Our CODEX data analysis relied on a simple featurization that collapsed single-cell images to mean intensities per channel, corresponding to protein abundances. To investigate the information contained in more complex, spatially resolved features, we turned to a joint spatial transcriptomics and fluorescence microscopy dataset of human ovarian cancer acquired using the 10x Genomics Xenium platform^57^ (Fig. 2a, b). Using the provided segmentation masks we extracted 1.6 million single-cell images across four channels with scPortrait (Fig. 2c). We aimed to embed these cells into a continuous representation space based on their morphologies alone. Rather than biasing this representation towards predefined labels, we adopted a self-supervised learning approach to retain as much cell morphology information as possible. This paradigm has been shown to generate meaningful image representations irrespective of external labels by training on auxiliary tasks such as reconstructing masked image patches^58^.

**Figure 2.**
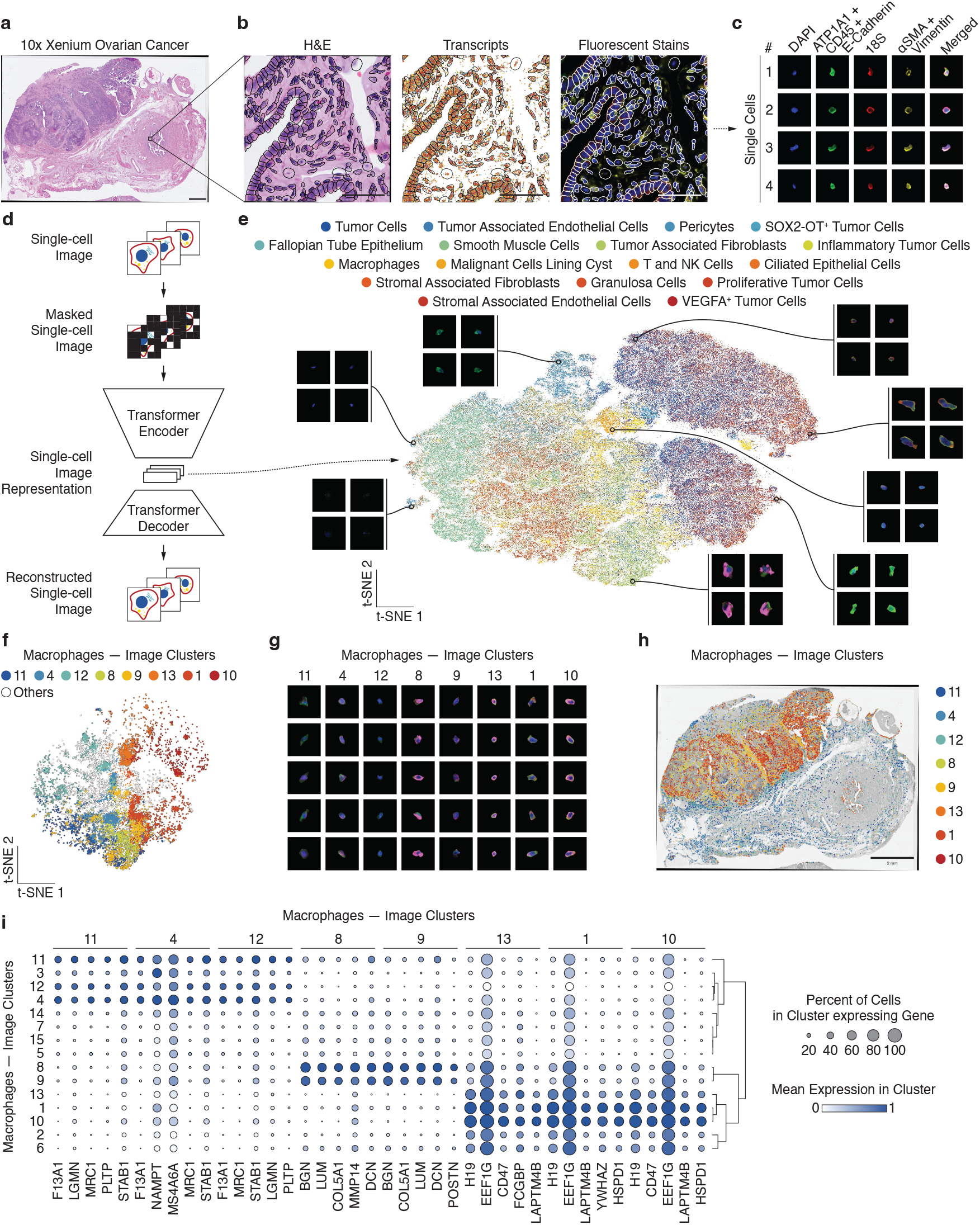
scPortrait identifies macrophage subpopulations in ovarian cancer. **a**, H&E overview image of tissue region contained in 10x Genomics Xenium dataset of human ovarian cancer. Scalebar represents 1 mm. **b**, Magnified region of a, black outlines show cell cytosol borders. Different panels show modalities contained in the dataset. Dots in center panel correspond to probes binding individual transcripts. Scalebars represent 50 µm. **c**, Single-cell images extracted with scPortrait. ATP1A + CD45 + E-Cadherin and αSMA + Vimentin were stained together in single imaging channels. We extracted a total of 1,627,204 single-cell images across four channels. The images were min-max scaled for visualization. **d**, Overview of single-cell image embedding strategy. We trained a transformer-based encoder-decoder model on the ATP1A + CD45 + E-Cadherin, 18S and αSMA + Vimentin channels via mask reconstruction (ViT-MAE) independently of biological labels. We used the internal representation learned by this model as an image-based featurization of cells in the dataset. **e**, t-SNE visualization of ViT-MAE embeddings of single-cell images colored by dataset cell type annotation. Each dot represents one cell. Images from indicated regions were not rescaled. **f**, e filtered for macrophages. Colors indicate selected Leiden clusters. **g**, Single-cell images of macrophages from clusters in f. **h**, Distribution of macrophage clusters from f across the tissue region. **i**, Genes differentially expressed in macrophage clusters from f.

We fine-tuned a vision transformer-based autoencoder (ViT-MAE)^59^ on the ATP1A/CD45/E-Cadherin, 18S and αSMA/Vimentin channels of the ovarian cancer dataset by mask reconstruction (Fig. 2d). The representation learned by this model’s encoder captured meaningful biology, as demonstrated by its ability to separate single-cell images by their morphological phenotypes in line with a previous report^60^ (Fig. 2e, S4a). Remarkably, it had also implicitly learned to group cells by type despite never being trained on cell type labels (Fig. 2e, S4a). To test whether this image-based representation contained information absent from the transcriptome, we inspected the difference in local neighborhood structure for each cell in both image- and transcriptome space. If the two modalities contained similar information, the same cells should be neighbors of one another in each embedding. Average neighborhood overlap was less than 5% across cell types, demonstrating that our image-based embedding encodes largely non-redundant information (Fig. S4b, c). Tumor cells generally exhibited the highest overlap while T and NK cells showed the lowest (Fig. S4c).

To probe the information captured by our fine-tuned ViT-MAE model in more detail, we focused on macrophages, a heterogeneous and functionally diverse cell type. Clustering all macrophages by image features revealed several morphologically distinct subclusters (Fig. 2f, g, S4d). Their spatial distributions differed strikingly, particularly in their intra-versus extratumoral localization (Fig. 2h, S4e, f). Thus, morphology alone was sufficient to distinguish macrophage states associated with distinct tissue niches, demonstrating that image-based embeddings can resolve biologically meaningful heterogeneity.

To explore the functional characteristics of these morphologically distinct macrophages, we turned to their spatial transcriptomics profiles. Differential gene expression analysis revealed cluster-specific signatures (Fig 2i, S4g): Clusters 4, 11 and 12, predominantly extra-tumoral, express *MRC1*, encoding CD206, and *STAB1*, markers of anti-inflammatory macrophage subpopulations^61,62^. In contrast, clusters 8 and 9 were almost exclusively found intratumoral and expressed *LUM* and *COL5A1*, consistent with a population of cancer-associated fibroblasts mislabeled as macrophages^63,64^. These results show that morphology-based clustering by representation learning with scPortrait can resolve functionally defined cell states from images.

### scPortrait embeds cells into a transcriptome atlas based on their images

Large, labeled single-cell collections are increasingly becoming available^65–67^, but most of these atlases are centered around single-cell transcriptomics. After using single-cell images to identify an anti-inflammatory macrophage subset, we asked whether single-cell images could be annotated directly from transcriptome atlases. We reasoned that a small amount of overlap in image- and transcriptome-based information might be sufficient to embed cells into transcriptomic atlases directly from their images.

We first projected all cells of the ovarian cancer data set into the SCimilarity atlas^65^ based on their transcriptome. Then, using the ViT-MAE image features we generated as inputs, we trained a multilayer perceptron (MLP) as a cross-modality model to predict SCimilarity features from the ViT-MAE image embeddings (Fig. 3a). For all analyses of these data we excluded tumor cells, since SCimilarity is not trained to properly embed them^65^. On a held-out test set this cross-modality model achieved an R^2^ value of 0.65 (Fig. 3b). Inspecting assigned ovary cell type labels showed that a variety of cell types was predicted, and similar cell types clustered together. To understand the biases of our model we then investigated the prediction error by cell type. Error analysis revealed similar performance across most cell types, with epithelial cells and T / NK cells showing the highest errors, suggesting that variance was not dominated by a single lineage (Fig. 3c).

**Figure 3.**
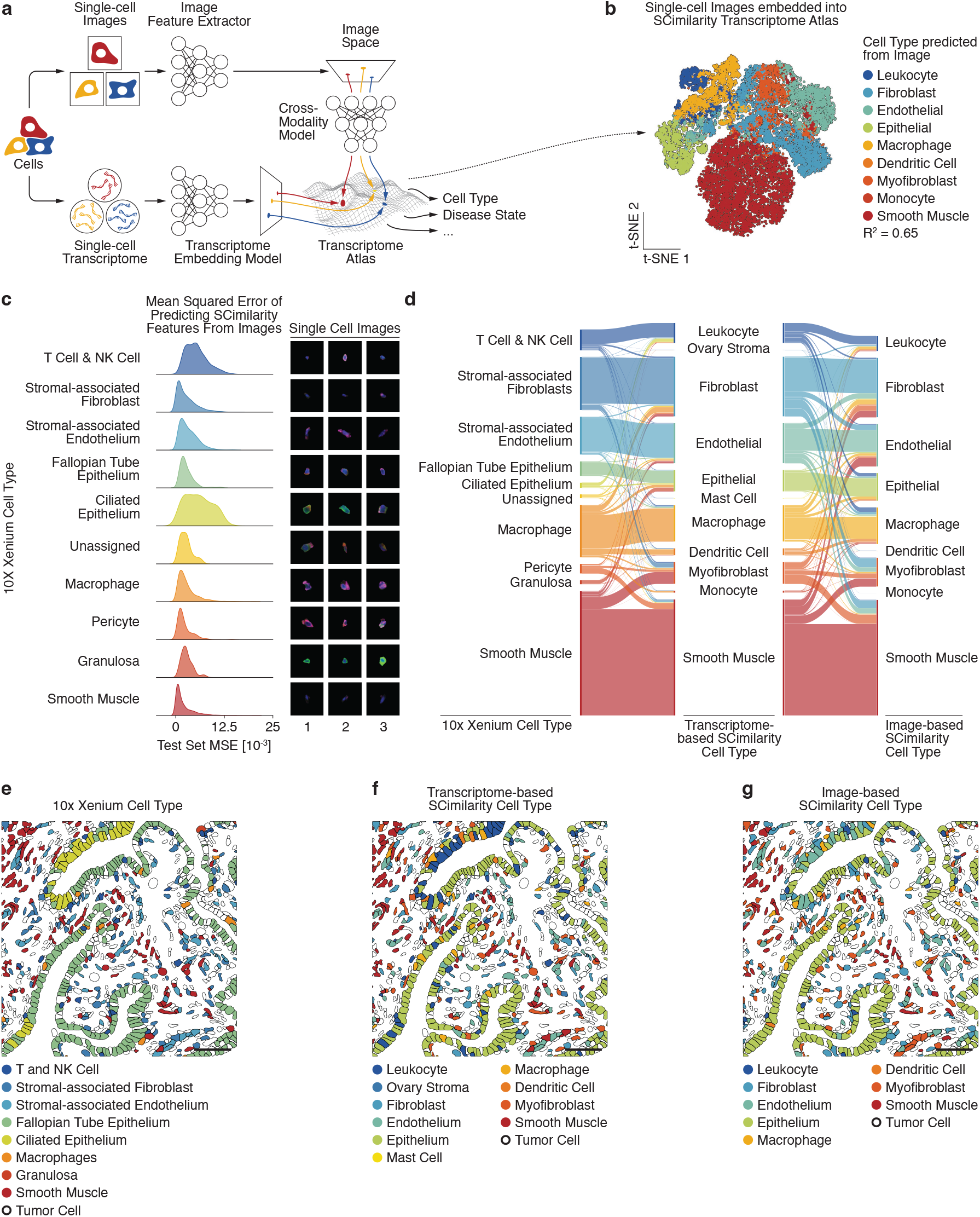
scPortrait enables cross-modality embedding of single-cell images into a transcriptome atlas. **a**, Overview of our strategy to embed single cells into a transcriptome atlas by their images. First, we embed all cells from the ovarian cancer dataset into the single-cell transcriptome atlas SCimilarity by their transcriptome. We then train a multilayer perceptron (MLP) model to predict a cell’s SCimilarity embedding from its ViT-MAE image embedding. This model embeds cells into SCimilarity even if only low-quality transcriptome information is available, based solely on their images. **b**, t-SNE of test set cells embedded into SCimilarity atlas, colored by SCimilarity cell type label. **c**, Left: Mean squared error (MSE) of SCimilarity embedding prediction from single-cell images by cell type. Only test set data are shown. Right: Corresponding single-cell images. **d**, Comparison of cell type labels in the ovarian cancer dataset (left) to transcriptome-derived SCimilarity embeddings (center) and image-derived SCimilarity embeddings (right). Only test set data are shown. **e, f, g**, Spatial distribution of cell type labels from the ovarian cancer dataset (e), transcriptome-derived SCimilarity embeddings (f) and image-derived SCimilarity embeddings (g). The depicted tissue region was excluded from the training set. Scalebars represent 50 µm.

An advantage of projecting cells of new experiments into existing atlases is that atlas labels can be used to infer biological information about previously unlabeled samples. Given that the ovarian cancer dataset contained cell type labelling already, we compared the provided cell types with those predicted by transcriptome- or image-based SCimilarity embeddings. Most cells showed strong agreement across all three labels (Fig. 3d, S5a, b) and mismatches typically involved related types such as smooth muscle cells and myofibroblasts. In line with the increased prediction error, ciliated epithelium and leukocytes were notable exceptions (Fig. 3c, d). To further validate the approach, we mapped predicted cell types onto a tissue region not used for training. Both transcriptome- and image-derived SCimilarity cell type labels annotated the tissue correctly, identifying epithelial structures, fibroblasts and smooth muscle cells (Fig. 3e, f, g). Together, these results demonstrate that cells can be embedded into transcriptome atlases based on images alone, recovering meaningful biological information and underscoring the role of imaging in multimodal integration and modeling.

## Discussion

Microscopy is one of the most information-rich and scalable ways to study cells, yet image data have lagged behind other single-cell modalities in standardization and integration. While transcriptomics and proteomics already rely on widely adopted formats that enable large-scale analysis, image-based datasets remain fragmented, often tied to specific pipelines or instruments. scPortrait addresses this gap by introducing .h5sc, a standardized and accessible format for single-cell images, together with a scalable software framework for dataset generation and sharing.

Open formats are crucial for modern life science research^47^. By building .h5sc on HDF5 and the AnnData specification^37^, scPortrait ensures compatibility with the scverse ecosystem and downstream tools such as squidpy^44^ or bento^40^. This design promotes interoperability, reproducibility, and reuse, aligning single-cell imaging with the FAIR principles^46^. In addition, .h5sc delivers fast random access to individual cells — essential for training large machine learning models — and avoids repeated, compute-intensive preprocessing. Thereby, scPortrait elevates single-cell imaging to the same level of accessibility and reuse that transcriptomics has already achieved, filling a critical community need.

A key advantage of images is the short turnaround time of their acquisition, enabling iterative evaluations with machine learning models in a loop. Paradigms such as active learning^68^, where new observations are specifically sampled to address a model’s current uncertainties, benefit from faster iteration times with scPortrait. Such approaches can be used in real time to regulate image acquisition parameters, enabling microscopes to dynamically respond to sample characteristics^69^. Across experiments, reinforcement learning can guide experimental design: A model would use the results from a given round of imaging experiments combined with the knowledge from already available representations learned by existing models^70^ to recommend a new set of experiments for the next round, iteratively exploring the underlying biology. Reinforcement learning benefits directly from consistent representation of data at a defined level, achieved through the standardized .h5sc single-cell image datasets created by scPortrait.

In this study, we showed how scPortrait enables analyses that go beyond current imaging workflows. Using self-supervised representations of ovarian cancer tissue, we identified macrophage subpopulations with distinct spatial distributions and transcriptomic signatures, illustrating how morphology can resolve biologically meaningful cell states. In tonsil tissue, cross-modality mapping and flow matching allowed us to infer gene expression directly from CODEX images, recovering missing markers such as TCL1A and revealing tissue organization without matched samples. Finally, embedding cells into the SCimilarity transcriptome atlas demonstrated that morphology-derived features can generalize across datasets, opening the door to image-driven atlas annotation.

Beyond its immediate applications, image-based modeling raises unique challenges that will shape the next stage of single-cell analysis. Imaging experiments vary widely in hardware, staining protocols and preprocessing, leading to strong batch effects that hinder integration across datasets^71.^ Similar challenges have been addressed in transcriptomics and proteomics^71,72,^ and a standardized format such as .h5sc can catalyze comparable solutions for imaging by making large, diverse datasets broadly accessible.

Another limitation is interpretability: image embeddings often lack clear biological meaning compared with transcript or protein abundances. Here, integration across modalities provides a powerful remedy. In our ovarian cancer analysis, scPortrait-derived morphology separated macrophage subsets whose molecular identities were clarified by transcriptomic profiles. Such cross-modality approaches not only validate image-based findings but also enable the discovery of new cell states that might otherwise remain hidden.

In conclusion, scPortrait provides the infrastructure needed to place imaging alongside transcriptomics and proteomics as a core modality for single-cell biology. By enabling large-scale sharing and integration, it supports the development of multimodal foundation models that learn from images as well as molecular profiles. Such models hold the promise of capturing complementary aspects of cell identity and behavior, ultimately enabling a more complete and predictive understanding of human biology.

## Methods

### The scPortrait software package

The Python-based scPortrait software package is available on GitHub (https://github.com/MannLabs/scPortrait) with documentation and tutorials (https://mannlabs.github.io/scPortrait/). scPortrait reads raw microscopy data and then follows a processing pipeline that generates standardized single-cell image datasets via three main steps as outlined here (https://mannlabs.github.io/scPortrait/pages/workflow.html):

- Stitching
- Segmentation
- Extraction

Two additional steps can follow:

- Featurization
- Selection of cells for downstream computational or biological analysis

All processing steps are carried out on an scPortrait project that defines a data structure on disk for saving intermediate and final results. One config file per project specifies options for each of the above steps. A variety of workflows are available for each step and can be used directly or adapted to suit specific dataset characteristics.

### Processing of human tonsil CODEX data

Raw TIFF images were percentile-normalized to the 0.1 % - 99.9 % range per channel and then separated by disease status into healthy and tonsillitis tissue cores. These images were then segmented with scPortrait using the CellPose^32^ nucleus model followed by mask expansion to generate cell borders. Single-cell images were then extracted into 36 × 36 px images. To preserve relative signal strengths across cells for each channel we did not rescale single-cell image intensities. Single cells were then featurized using the scPortrait CellFeaturizer, and mean intensity per channel was used for downstream analyses.

### Mapping CODEX features to single-cell RNA-seq with generative Optimal Transport

#### Problem statement

Given a dataset of *N* CODEX cell images represented by protein marker features measured by scPortrait, our goal is to derive an approximation of the gene expression profiles of the imaged cells, leveraging a multi-modal single-cell reference dataset containing gene expression counts and protein markers.

Formally, let *Y* ∈ ℝ^*N x D*^ be the matrix of *N* imaged cells across *D* marker-specific features. Moreover, denote *X*^*p*^ ∈ ℝ^*M x D*^ and *X*^*g*^ ∈ ℝ^*M x G*^ the *cell × protein* and *cell × gene* matrices in a reference single-cell CITE-seq atlas from the same tissue as the CODEX samples. Note that *X*^*p*^ has the same number of features as *Y*, as we subset the two feature spaces to reflect the same measured protein markers. Our goal is to derive a predicted gene expression matrix 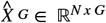 for each cell imaged by CODEX.

Our approach consists of two steps:

1. Match CODEX samples to their putative single-cell counterpart based on the similarity between image-based and CITE-seq-derived protein marker abundances. We pair the two modalities with Optimal Transport (OT).
2. Learn a mapping that transports CODEX features to the single-cell gene expression measured in the atlas.

To account for the noise in the single-cell dataset, we train a generative model based on flow matching^29,30^ that generates novel gene expression profiles using the image-based marker features as input.

#### Learning OT with flow matching

Flow matching learns a parameterized, time-resolved vector field *v*^*θ*^_*t*_(*x*) that maps samples from a prior distribution *N* (0, *I*) (by convention, at *t* = 0) to a target data distribution *p* at *t* = 1, thereby acting as a generative model by turning noise into data samples via the following equation:

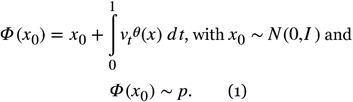

The integral function *Φ* mapping noise to data is called *flow*.

Klein et al.^28^ showed that one can train flow matching to approximate a generative OT map from a source distribution *q* to the target data distribution *p* described above by conditioning the generative process with samples *y* ~ *q*from the source:

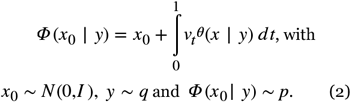

In other words, flow matching learns to transport a sample *y* from the source to the target stochastically according to an OT criterion, based on a pre-defined cost function.

#### Model training

In our setting, where the aim is to map cells from single-cell images in CODEX to single-cell CITE-seq, we indicate the image-based feature distribution as *q* and the single-cell CITE-seq distribution as *p*. We parameterize the velocity function *v*_*t*_^*θ*^(*x*) using a neural network trained with stochastic gradient descent over minibatches. One training iteration consists of the following steps:

1. Draw a random batch of *B* samples *Y*_*b*_ = {*y*_*i*_}^*B*^_*i*=1_ from the CODEX dataset
2. Draw a random batch of *B* single cells from the multi-modal reference atlas in both their protein and gene expression views as a target. We denote the target batch with *X*_*b*_ = (*X*_*b*_^*G*^, *X*_*b*_^*P*^) = {(*x*^*G*^_*i*_, *x*^*P*^_*i*_)}^*B*^_*i*=1_.
3. Compute an OT coupling matrix *Π* between all observations in *Y*_*b*_ and *X*_*b*_^*P*^ minimizing the following *Euclidean cost: c*(*y*_*i*_, *x*^*P*^_*i*_) = | | *y*_*i*_ − *x*_*i*_^*P*^ | |_2_. The coupling approximates a joint distribution between source and target sample indices, where CODEX cells are mapped with a higher probability to atlas cells with similar marker abundance.
4. Resample couples of source and target indices from the joint distribution, yielding the resampled batches 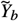 and 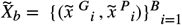. The *i*^*th*^ sample in 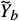 is transported to the *i*^*th*^ sample in 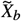 with high probability *Π*.
5. Perform a flow matching iteration^30^, training the vector field to transport a noise sample *x*_0_ ~ *N* (0, *I*) to the gene expression vector 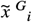 conditioned on its matched CODEX feature vector 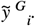.

#### Inference

During inference, we transport a CODEX sample *y* to its gene expression counterpart by sampling a noise point *x*_0_ ∈ *N* (0,*I*) and computing 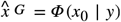.

#### Data preprocessing

We subset the CODEX features extracted by scPortrait to the mean intensity of the marker channels and standardize individual features to mitigate skewness towards zero. To ensure correspondence between image and single-cell protein abundance data, we subset the CODEX features extracted by scPortrait and the single-cell protein abundance measurements to their 31 shared markers.

Since flow matching is a model working in continuous space, we use batch-corrected and 50-dimensional expression features extracted by the HARMONY algorithm^73^ as a target representation for generation. To predict gene expression from translated CODEX features, we pre-train a decoder function that maps HARMONY embeddings to gene expression.

#### Model architecture and training details

1. The flow matching velocity model *v*^*θ*^_*t*_(*x*) is an MLP with 3 layers of 1024 hidden dimensions each and an ELU activation function. We train it with the AdamW optimizer with a learning rate of 1e-4, a batch size of 256 and across 2000 epochs. The velocity function *v*^*θ*^_*t*_(*x*) inputs a concatenation of the current state with a generation time embedding. To embed the generation time, we use a sinusoidal encoding^1^ with 128 dimensions.
2. The decoder function, which maps HARMONY features to normalized gene expression, is an MLP with two layers of 64 hidden dimensions. We train the model for 200 epochs with a batch size of 256 and a learning rate of 1e-3. We choose a mean-squared error (MSE) loss function to reconstruct gene expression.

### Processing of human ovarian cancer dataset

We downloaded the Xenium ovarian cancer dataset from 10x Genomics using this link: https://www.10xgenomics.com/datasets/xenium-prime-ffpe-human-ovarian-cancer. After reading the dataset into SpatialData^35^, we generated single-cell images with scPortrait by extracting into 224 × 224 px images using the segmentation masks provided with the dataset. To preserve relative signal strengths across cells for each channel we did not rescale image single-cell intensities.

### Golgi morphology experiments

On day 0, 1 million HeLa cells expressing hsTGOLN2mCherry were plated per well of a 4-well plate containing a UV-sterilized metal frame slide with a polyphenylene sulfate (PPS) membrane. Cells were cultured in DMEM supplemented with 10 % FCS, 1 mM pyruvate, 100 U/ml penicillin and 100 µg/mL streptomycin at 37 °C and 5 % CO^2^. On day 2, cells were treated with 10 µM Golgicide A, 10 µM Nigericin or 20 µM Monensin for 2 hrs or with 5 mg/mL Nocodazole for 30 minutes. Cells were then stained with 10 µg/mL WGA-Alexa-647 in PBS for 10 minutes at 37 °C before being washed 3× in PBS and then fixed in 4 % paraformaldehyde (PFA) diluted in PBS for 10 minutes at room temperature. After fixation, cells were washed 3× in PBS again before being stained with 10 µg/mL Hoechst 33342 for 15 minutes at room temperature. Afterwards, cells were washed 3× in PBS again and imaged on a Nikon Eclipse Ti2 spinning disc confocal microscope.

### Training ViT-MAE

We fine-tuned a vision transformer-based masked autoencoder (ViT-MAE^59^) on the ovarian cancer dataset to learn a representation of cell morphology in an unsupervised manner, starting from a model pre-trained on natural images (https://huggingface.co/facebook/vitmae-base)^74^. The raw images contained 4 stains: DAPI, ATP1A/CD45/E-Cadherin, 18S and αSMA/Vimentin. To match the input dimensions of the pretrained model we subsetted to three channels, discarding DAPI to focus on the functional structures of cells. The dataset contains 406,875 images of single cells, which were split into 90 %, 5 % and 5 % for training, validation and test sets respectively, including a spatially defined region in the test set. Prior to training, the images were cropped around the center at a fixed size of 128 × 128 and resized to 224 × 224 to match the expected ViT-MAE input size. The resulting single-cell image dimension was 3 × 224 × 224. No other augmentations were applied. The model was trained for 119 epochs using the scPortrait PyTorch dataloader with a batch size of 128^75^. After splitting the input image into 16 × 16 px patches, the encoder of the model passes the input through 12 transformer layer blocks, each of which uses multihead self-attention with 768 embedding dimensions per head and 12 attention heads. The patch size is 16 × 16. GELU is used as an activation function with a layernorm of eps=1e-12. The decoder of the network is lighter than the encoder but contains the same structure with 8 attention blocks of 512 embedding dimensions and 16 attention heads. The intermediate sizes of the feedforward layers are 3072 and 2048 for the encoder and the decoder. We use a masking ratio of 75 % of patches, which is set to 0 during inference after training. In total, the model has 111 million trainable parameters. We construct the latent space by average pooling across all tokens. After scaling to zero mean and unit variance we used the Leiden algorithm with a resolution of 0.25 for clustering the latent space on 15 neighbors.

### Embedding cells into SCimilarity atlas

To embed cells from the ovarian cancer dataset into SCimilarity, transcriptomics data were preprocessed as described in the SCimilarity documentation: Each cell was normalized to 10,000 counts, and the expression matrix was aligned with the SCimilarity atlas. Of note, we did not log1p transform our data, since the distribution of probe-based spatial transcriptomics counts is different from stochastically sampled dissociated transcriptomics assays. We then calculated SCimilarity features for all cells in the dataset. The SCimilarity checkpoint used was downloaded from https://zenodo.org/records/10685499.

To embed cells into SCimilarity based on their images, we trained a multilayer perceptron (MLP) to predict transcriptome-derived SCimilarity features from per-vector min-max scaled ViT-MAE embeddings. The MLP consisted of five linear layers gradually decreasing in size from the 768 dimensions of the ViT-MAE features to the 128 dimensions of the SCimilarity features. ReLU was used as an activation function. The model was trained to minimize the mean squared error (MSE) of predicted to transcriptome-derived SCimilarity features with a learning rate of 1e-5. Model training was done in PyTorch Lightning^75,76^, with minimal validation loss as a selection criterion for the final set of model parameters used during inference. The same test set was held out from both ViT-MAE training and cross-modality MLP training, including a spatially defined region.

When predicting cell types for the ovarian cancer dataset we limited the list of possible cell types to those typically found in the ovary. All tumor cells were excluded from SCimilarity cell type predictions, because tumor cells were not part of the training data.

### Differential marker expression testing

Subgroups for testing of marker enrichment were assigned by Leiden clustering in scanpy or via external labels such as cell types. Differentially enriched markers or expressed genes were determined using scanpy’s *rank_genes_groups* function. The results were visualized using *rank_genes_groups_dotplot*.

## Data Availability

All data generated as part of this work are available at https://zenodo.org/records/17162225 or can be regenerated from publicly available sources using the code in the repository at the following URL: https://github.com/MannLabs/scPortrait_manuscript.

## Code Availability

The scPortrait software package is available at https://github.com/MannLabs/scPortrait. All figures in this manuscript can be recreated using the code in the repository at the following URL: https://github.com/MannLabs/scPortrait_manuscript.

## Acknowledgements

We would like to thank the Center for Advanced Light Microscopy (CALM) for support with light microscopy; Marvin Thielert and Marc Oeller for testing new features of scPortrait; Magnus Schwörer for helping automate scPortrait’s GitHub workflows; Piero Coronica from the Max Planck Computing and Data Facility (MPCDF) for support with improving the memory footprint of scPortrait’s single-cell image dataset class during model training; and Ilan Gold for discussions about strategies for developing scPortrait as an scverse package from the ground up. We thank all scPortrait contributors and users who provided feedback. We are grateful to the communities behind the multiple open-source software packages on which we depend.

## Author Contributions

S.C.M., N.A.S., F.J.T. and M.M. conceived the study. S.C.M., N.A.S., V.H., F.J.T. and M.M. designed experiments. S.C.M., N.A.S., A.N., L.H., G.W., F.J.T. and M.M. conceived the scPortrait software. S.C.M., N.A.S., A.P., A.N., A.O.C., V.V., M.A. and G.W. performed experiments. S.C.M., N.A.S., A.P., V.H., F.J.T. and M.M. wrote the manuscript with input from all authors.

## Supplementary Figures

**Figure S1.**
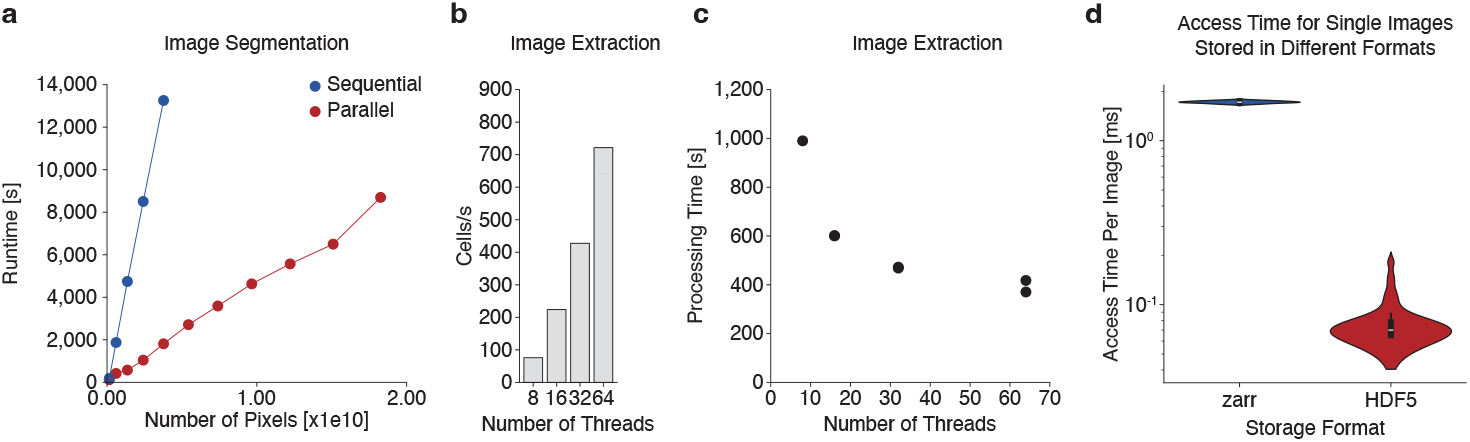
The parallelization capabilities of scPortrait and its HDF5 backend enables fast image processing and access. **a**, Runtime comparison of parallel and sequential image segmentation. **b**, Image extraction speed in cells per second over different numbers of concurrent threads. **c**, Image extraction processing time over different numbers of concurrent threads. **d**, Access duration of image datasets locally stored as HDF5 or zarr files. 100 images were accessed per file. s = seconds

**Figure S2.**
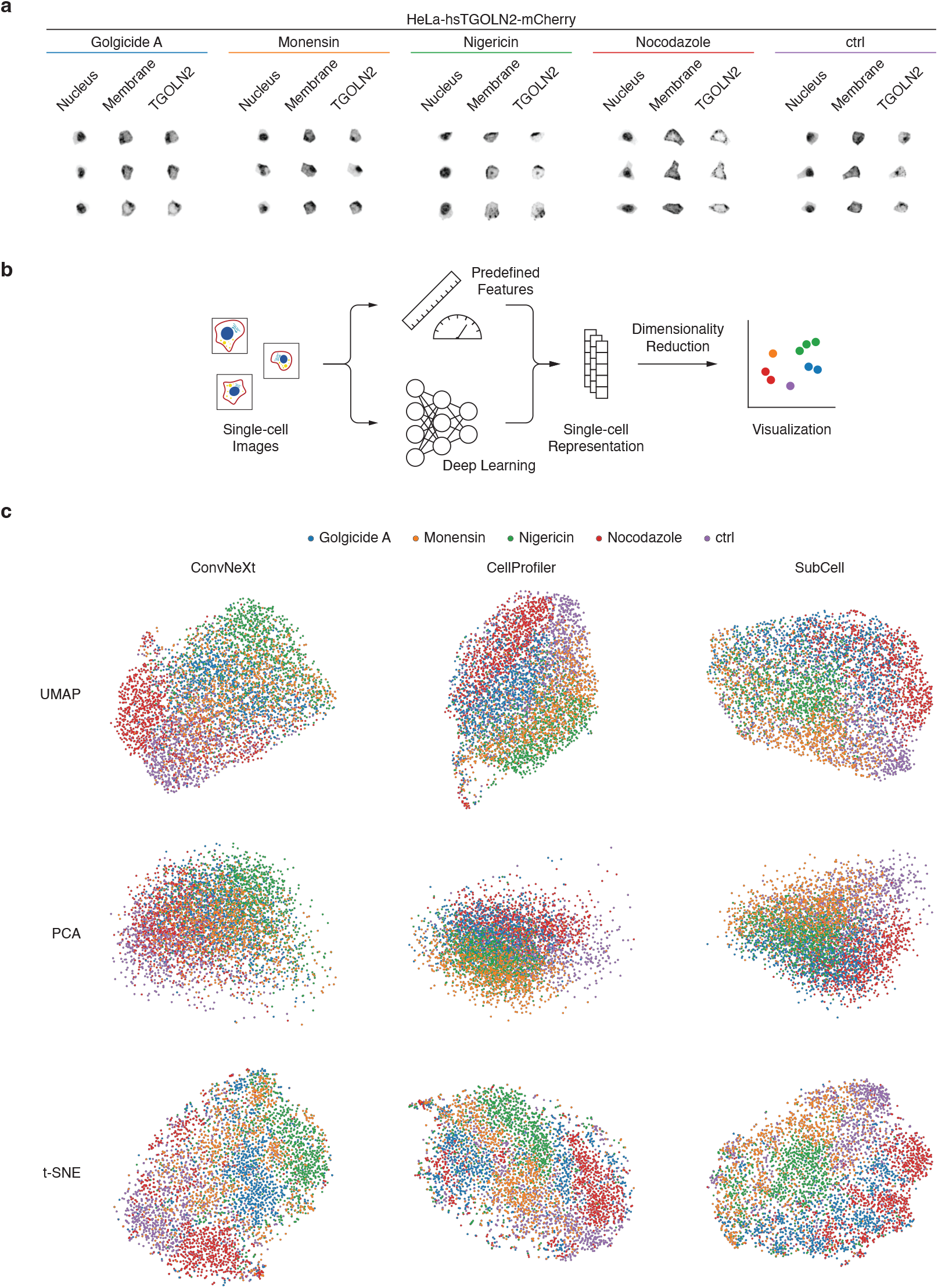
scPortrait recognizes trans-Golgi network morphologies in combination with different single-cell image embedding strategies. **a**, Single-cell images of HeLa cells expressing hsTGOLN2-mCherry stimulated with the indicated compounds. **b**, Embedding strategies for images from a. **c**, Three different single-cell image embedders were used to generate representations of images shown in a: ConvNeXt, a CNN trained as a classifier on natural images, CellProfiler, a list of predefined single-cell image features and SubCell, a transformer model trained on human protein atlas images. The representations learned by these models are visualized via three different techniques: t-SNE, UMAP and PCA. Colors indicate chemical perturbations from a.

**Figure S3.**
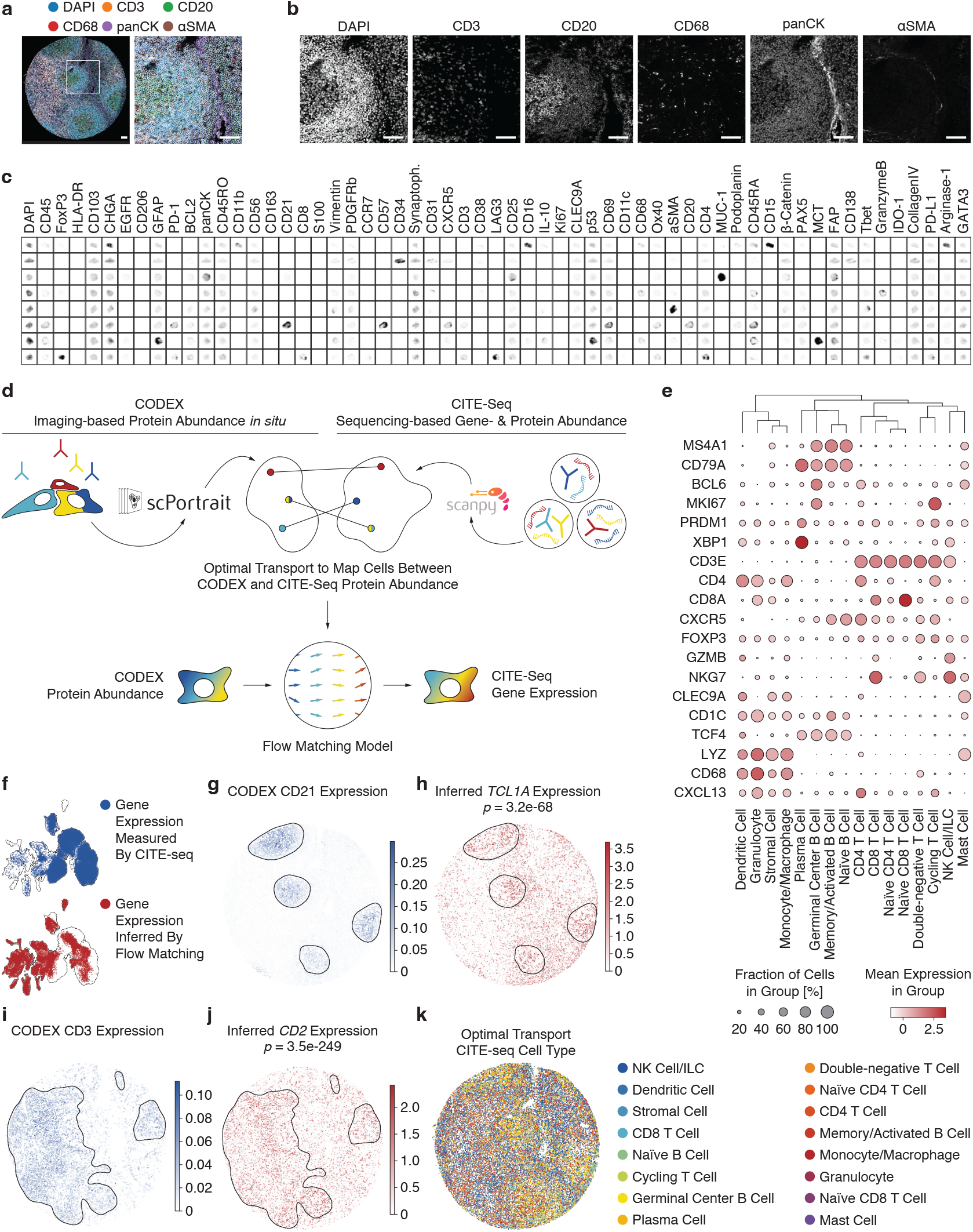
Optimal Transport matches CODEX imaging of human tonsil with transcriptome data. **a**, Overview of human tonsil tissue core from a tonsillitis patient stained with 58 antibodies and DAPI using the CODEX assay. Right panels depict magnification of regions indicated on the left. Colors depict selected stains. White outlines indicate cytosol borders determined by nuclear expansion segmentation based on CellPose. Scalebars represent 75 µm. **b**, Individual imaging channels of magnified region from a. Scalebars represent 75 µm. **c**, Example single-cell images following scPortrait extraction showing all 59 channels per cell. “Synaptoph.” = Synaptophysin. **d**, Overview of our strategy to match image-based CODEX data of human tonsil with unpaired single-cell transcriptomics data of human tonsil using optimal transport to train a flow matching model. We process the respective data modalities with scPortrait (images) and scanpy (transcriptomics), collapsing images to median channel intensity per cell. We then calculate a probabilistic mapping between modalities using Monge optimal transport. Sampling batches according to this optimal transport coupling between modalities, we then train a flow matching model to generate a cell’s gene expressions conditioned on its CODEX profile. **e**, Flow matching-inferred expression of selected marker genes across CITE-seq derived cell types. Cell types were assigned to inferred expression profiles by k-nearest neighbor majority. **f**, UMAP representation of gene expression measured by CITE-seq or inferred by flow matching per cell. Outlines correspond to joint UMAP. **g**, Germinal center marker CD21 expression measured by CODEX in the tonsillitis sample. Black outlines show germinal centers defined by the expression of this marker. **h**, *TCL1A* expression inferred by flow matching in the tonsillitis sample. Black outlines show germinal centers defined by the expression of CD21. **i**, T cell marker CD3 expression measured by CODEX in the tonsillitis sample. Black outlines show T cell zone borders defined by the expression of this marker. **j**, *CD2* expression inferred by flow matching in the tonsillitis sample. Black outlines show T cell zone borders defined by the expression of CD3. **k**, Cell type annotation from dissociated transcriptomics data mapped onto tonsillitis tissue via optimal transport. *p*-values correspond to spatially differential gene expression in- and outside of the highlighted zones and were calculated by a two-sided Mann-Whitney U test with Bonferroni correction.

**Figure S4.**
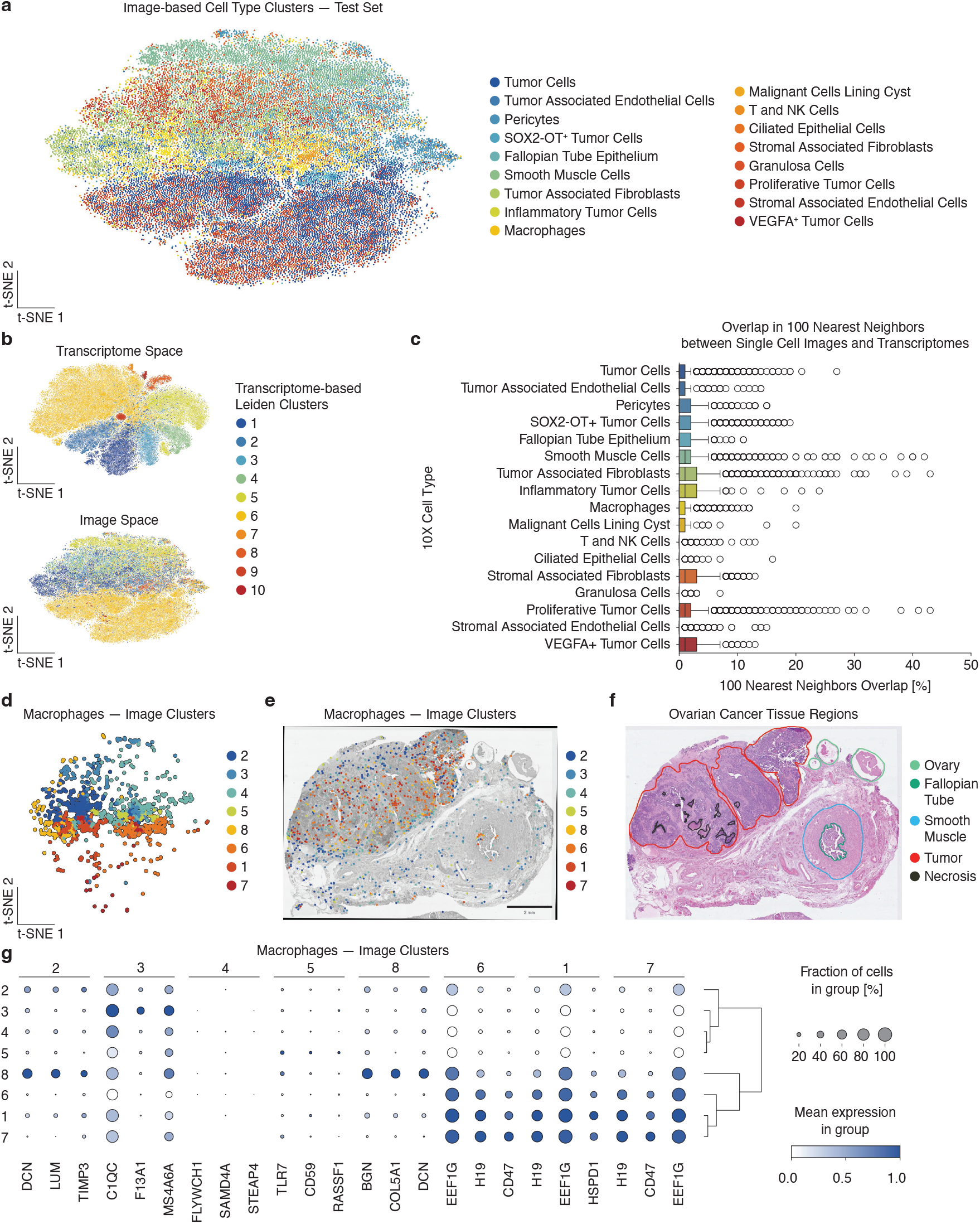
Cross-modality modeling of spatial transcriptomics and imaging data with scPortrait. **a**, t-SNE visualization of ViT-MAE embeddings of single-cell images colored by 10x Xenium cell type annotation. Each dot represents one cell. Only test set data are shown. **b**, t-SNE visualization of single-cell transcriptome (top) and ViT-MAE embeddings of single-cell images (bottom) colored by transcriptome-based Leiden clusters. Each dot represents one cell. Only test set data are shown. **c**, Boxplot depicting local neighborhood overlap between transcriptome space and image-based embeddings (b). X-axis shows percent of neighbors of a given cell that are identical between transcriptome space and image-based embeddings. Y-axis shows 10x cell types. **d**, a filtered for macrophages. Colors indicate Leiden clusters. Only test set data are shown. **e**, Distribution of macrophage clusters from d across the tissue region. Only test set data are shown. **f**, Tissue region annotation of ovarian cancer dataset. **g**, Genes differentially expressed in macrophage clusters from d. Only test set data are shown.

**Figure S5.**
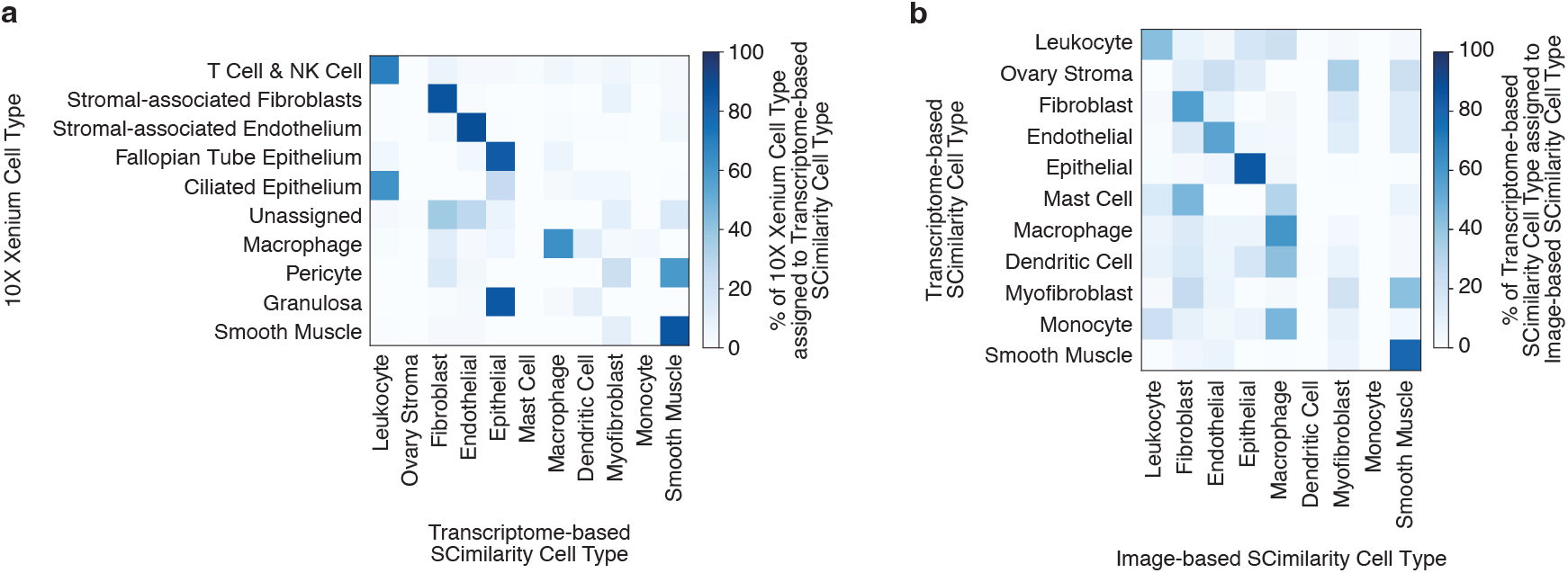
Embedding of single-cell images into a transcriptome atlas with scPortrait. **a**, Heatmap showing which cell types cells get assigned after embedding into SCimilarity space based on their transcriptome as a percentage of 10x Xenium cell types. **b**, Heatmap showing which cell types cells get assigned after embedding into SCimilarity space based on their images as a percentage of transcriptome-based SCimilarity cell types.

## References

1. Vaswani, A. et al. Attention is all you need. Advances in neural information processing systems 30 (2017).

2. LeCun, Y., Bengio, Y. & Hinton, G. Deep learning. Nature 521, 436–444 (2015).

3. Schmidhuber, J. Deep Learning in Neural Networks: An Overview. arXiv, 1404.7828 (2014).

4. Krizhevsky, A., Sutskever, I. & Hinton, G. E. ImageNet Classification with Deep Convolutional Neural Networks. Advances in neural information processing systems 25 (2012).

5. Price, I. et al. Probabilistic weather forecasting with machine learning. Nature 637, 84–90 (2025).

6. Bunne, C. et al. How to build the virtual cell with artificial intelligence: Priorities and opportunities. Cell 187, 7045–7063 (2024).

7. Yirmiya, E. et al. Structure-guided discovery of viral proteins that inhibit host immunity. Cell 188, 1681–1692.e1617 (2025).

8. Seal, S. et al. Cell Painting: a decade of discovery and innovation in cellular imaging. Nature Methods 22, 254–268 (2025).

9. Thul, P. J. et al. A subcellular map of the human proteome. Science 356, eaal3321 (2017).

10. Chung, K. et al. Structural and molecular interrogation of intact biological systems. Nature 497, 332–337 (2013).

11. Bae, J. A. et al. Functional connectomics spanning multiple areas of mouse visual cortex. Nature 640, 435–447 (2025).

12. Chandrasekaran, S. N. et al. Three million images and morphological profiles of cells treated with matched chemical and genetic perturbations. Nat Methods 21, 1114–1121 (2024).

13. Bock, C. et al. High-content CRISPR screening. Nature Reviews Methods Primers 2, 8 (2022).

14. Schmacke, N. A. et al. SPARCS, a platform for genome-scale CRISPR screening for spatial cellular phenotypes. bioRxiv, 2023.2006.2001.542416 (2023).

15. Pitino, E. et al. STAMP: Single-cell transcriptomics analysis and multimodal profiling through imaging. Cell 188, 5100–5117.e5126 (2025).

16. Mund, A. et al. Deep Visual Proteomics defines single-cell identity and heterogeneity. Nature Biotechnology 40, 1231–1240 (2022).

17. Tian, L., Chen, F. & Macosko, E. Z. The expanding vistas of spatial transcriptomics. Nature Biotechnology 41, 773–782 (2023).

18. Cui, H. et al. Towards multimodal foundation models in molecular cell biology. Nature 640, 623–633 (2025).

19. Ji, Y. et al. Scalable and universal prediction of cellular phenotypes. bioRxiv, 2024.2008.2012.607533 (2024).

20. Bussi, Y. & Keren, L. Multiplexed image analysis: what have we achieved and where are we headed? Nature Methods 21, 2212–2215 (2024).

21. Tan, Y. et al. SPACEc: A Streamlined, Interactive Python Workflow for Multiplexed Image Processing and Analysis. bioRxiv, 2024.2006.2029.601349 (2024).

22. Schapiro, D. et al. MCMICRO: a scalable, modular imageprocessing pipeline for multiplexed tissue imaging. Nature Methods 19, 311–315 (2022).

23. Kuehl, M. et al. Pathology-oriented multiplexing enables integrative disease mapping. Nature 644, 516–526 (2025).

24. Bankhead, P. et al. QuPath: Open source software for digital pathology image analysis. Scientific Reports 7, 16878 (2017).

25. Stirling, D. R. et al. CellProfiler 4: improvements in speed, utility and usability. BMC Bioinformatics 22, 433 (2021).

26. Muñoz, A. F. et al. cp_measure: API-first feature extraction for image-based profiling workflows. arXiv, 2507.01163 (2025).

27. Moore, J. et al. OME-NGFF: a next-generation file format for expanding bioimaging data-access strategies. Nature Methods 18, 1496–1498 (2021).

28. Klein, D., Uscidda, T., Theis, F. & Cuturi, M. GENOT: Entropic (Gromov) Wasserstein Flow Matching with Applications to Single-Cell Genomics. arXiv, 2310.09254 (2023).

29. Lipman, Y., Chen, R. T. Q., Ben-Hamu, H., Nickel, M. & Le, M. Flow Matching for Generative Modeling. arXiv, 2210.02747 (2022).

30. Tong, A. et al. Improving and generalizing flow-based generative models with minibatch optimal transport. arXiv, 2302.00482 (2023).

31. Muhlich, J. L. et al. Stitching and registering highly multiplexed whole-slide images of tissues and tumors using ASHLAR. Bioinformatics 38, 4613–4621 (2022).

32. Stringer, C., Wang, T., Michaelos, M. & Pachitariu, M. Cellpose: a generalist algorithm for cellular segmentation. Nature Methods 18, 100–106 (2021).

33. Mölder, F. et al. Sustainable data analysis with Snakemake [version 1; peer review: 1 approved, 1 approved with reservations]. F1000Research 10 (2021).

34. Di Tommaso, P. et al. Nextflow enables reproducible computational workflows. Nature Biotechnology 35, 316–319 (2017).

35. Marconato, L. et al. SpatialData: an open and universal data framework for spatial omics. Nature Methods 22, 58–62 (2025).

36. Sofroniew, N. et al. napari: a multi-dimensional image viewer for Python.

37. Virshup, I., Rybakov, S., Theis, F. J., Angerer, P. & Wolf, F. A. anndata: Access and store annotated data matrices. JOSS (2024).

38. Virshup, I. et al. The scverse project provides a computational ecosystem for single-cell omics data analysis. Nature Biotechnology 41, 604–606 (2023).

39. Hao, Y. et al. Dictionary learning for integrative, multimodal and scalable single-cell analysis. Nature Biotechnology 42, 293–304 (2024).

40. Mah, C. K. et al. Bento: a toolkit for subcellular analysis of spatial transcriptomics data. Genome Biology 25, 82 (2024).

41. Serrano, E. et al. Reproducible image-based profiling with Pycytominer. Nature Methods 22, 677–680 (2025).

42. Klein, D. et al. Mapping cells through time and space with moscot. Nature 638, 1065–1075 (2025).

43. Heumos, L. et al. Pertpy: an end-to-end framework for perturbation analysis. bioRxiv, 2024.2008.2004.606516 (2024).

44. Palla, G. et al. Squidpy: a scalable framework for spatial omics analysis. Nature Methods 19, 171–178 (2022).

45. Gayoso, A. et al. A Python library for probabilistic analysis of single-cell omics data. Nature Biotechnology 40, 163–166 (2022).

46. Wilkinson, M. D. et al. The FAIR Guiding Principles for scientific data management and stewardship. Scientific Data 3, 160018 (2016).

47. Bajcsy, P. et al. Enabling global image data sharing in the life sciences. Nature Methods 22, 672–676 (2025).

48. Liu, Z. et al. A ConvNet for the 2020s. arXiv, 2201.03545 (2022).

49. Gupta, A. et al. SubCell: Vision foundation models for microscopy capture single-cell biology. bioRxiv, 2024.2012.2006.627299 (2024).

50. Kim, V. et al. Self-supervision advances morphological profiling by unlocking powerful image representations. Scientific Reports 15, 4876 (2025).

51. Rosenberger, F. A. et al. Deep Visual Proteomics maps proteotoxicity in a genetic liver disease. Nature 642, 484–491 (2025).

52. Massoni-Badosa, R. et al. An atlas of cells in the human tonsil. Immunity 57, 379–399.e318 (2024).

53. Schiebinger, G. et al. Optimal-Transport Analysis of Single-Cell Gene Expression Identifies Developmental Trajectories in Reprogramming. Cell 176, 928–943.e922 (2019).

54. Klein, D. et al. CellFlow enables generative single-cell phenotype modeling with flow matching. bioRxiv, 2025.2004.2011.648220 (2025).

55. Huang, T., Liu, T., Babadi, M., Jin, W. & Ying, R. Scalable Generation of Spatial Transcriptomics from Histology Images via Whole-Slide Flow Matching. arXiv, 2506.05361 (2025).

56. Haviv, D., Pooladian, A.-A., Pe’er, D. & Amos, B. Wasserstein Flow Matching: Generative modeling over families of distributions. arXiv, 2411.00698 (2024).

57. 10x Genomics. Xenium Prime 5K In Situ Gene Expression with Cell Segmentation data for human ovarian cancer (FFPE) using the Xenium Prime 5K Human Pan Tissue and Pathways Panel plus 100 Custom Genes (v1), In Situ Gene Expression dataset analyzed using Xenium Onboard Analysis 3.0.0. (2024).

58. Giakoumoglou, N., Stathaki, T. & Gkelias, A. A Review on Discriminative Self-supervised Learning Methods in Computer Vision. arXiv, 2405.04969 (2024).

59. He, K. et al. Masked Autoencoders Are Scalable Vision Learners. arXiv, 2111.06377 (2021).

60. Kraus, O. et al. Masked Autoencoders for Microscopy are Scalable Learners of Cellular Biology. arXiv, 2404.10242 (2024).

61. Gordon, S. & Martinez, F. O. Alternative activation of macrophages: mechanism and functions. Immunity 32, 593–604 (2010).

62. Hollmén, M., Figueiredo, C. R. & Jalkanen, S. New tools to prevent cancer growth and spread: a ‘Clever’ approach. British Journal of Cancer 123, 501–509 (2020).

63. Muhl, L. et al. Single-cell analysis uncovers fibroblast heterogeneity and criteria for fibroblast and mural cell identification and discrimination. Nature Communications 11, 3953 (2020).

64. Kalluri, R. The biology and function of fibroblasts in cancer. Nature Reviews Cancer 16, 582–598 (2016).

65. Heimberg, G. et al. A cell atlas foundation model for scalable search of similar human cells. Nature 638, 1085–1094 (2025).

66. Mathys, H. et al. Single-cell multiregion dissection of Alzheimer’s disease. Nature 632, 858–868 (2024).

67. Sikkema, L. et al. An integrated cell atlas of the lung in health and disease. Nature Medicine 29, 1563–1577 (2023).

68. Huang, K. et al. Sequential Optimal Experimental Design of Perturbation Screens Guided by Multi-modal Priors. bioRxiv, 2023.2012.2012.571389 (2023).

69. Mahecic, D. et al. Event-driven acquisition for content-enriched microscopy. Nat Methods 19, 1262–1267 (2022).

70. Guo, D. et al. DeepSeek-R1 incentivizes reasoning in LLMs through reinforcement learning. Nature 645, 633–638 (2025).

71. Arevalo, J. et al. Evaluating batch correction methods for image-based cell profiling. Nat Commun 15, 6516 (2024).

72. Luecken, M. D. et al. Benchmarking atlas-level data integration in single-cell genomics. Nature Methods 19, 41–50 (2022).

73. Korsunsky, I. et al. Fast, sensitive and accurate integration of single-cell data with Harmony. Nature Methods 16, 1289–1296 (2019).

74. Wolf, T. et al. HuggingFace’s transformers: State-of-the-art natural language processing. arXiv (2019).

75. Falcon, W. & The PyTorch Lightning team. PyTorch Lightning, https://www.pytorchlightning.ai (2019).

76. Paszke, A. et al. PyTorch: An Imperative Style, High-Performance Deep Learning Library. arXiv, 1912.01703 (2019).

